# A unique form of collective epithelial migration is crucial for tissue fusion in the secondary palate and can overcome loss of epithelial apoptosis

**DOI:** 10.1101/2021.09.07.459343

**Authors:** Teng Teng, Camilla Teng, Vesa Kaartinen, Jeffrey O. Bush

## Abstract

Tissue fusion is an oft-employed process in morphogenesis which often requires the removal of the epithelia intervening multiple distinct primordia to form one continuous structure. In the mammalian secondary palate, a midline epithelial seam (MES) forms between two palatal shelves and must be removed to allow mesenchymal confluence. Abundant apoptosis and cell extrusion in this epithelial seam support their importance in its removal. However, by genetically disrupting the intrinsic apoptotic regulators BAX and BAK within the MES, we find a complete loss of cell death and cell extrusion, but successful removal of the MES, indicating that developmental compensation enables fusion. Novel static and live imaging approaches reveal that the MES is removed through a unique form of collective epithelial cell migration in which epithelial trails and islands stream through the mesenchyme to reach the oral and nasal epithelial surfaces. These epithelial trails and islands begin to express periderm markers while retaining expression of the basal epithelial marker ΔNp63, suggesting their migration to the oral and nasal surface is concomitant with their differentiation to an epithelial intermediate. Live imaging reveals anisotropic actomyosin contractility within epithelial trails that drives their peristaltic movement, and genetic loss of non-muscle myosin IIA-mediated actomyosin contractility results in dispersion of epithelial collectives and dramatic failure of normal MES migration. These findings demonstrate redundancy between cellular mechanisms of morphogenesis and reveal a crucial role for a unique form of collective epithelial migration during tissue fusion.

## Introduction

Tissue fusion is a complex morphogenetic process that occurs in diverse developmental contexts and organisms, involving the coordination of numerous cellular behaviors (Hashimoto et al., 2015; Hayes and Solon, 2017; Ray and Niswander, 2012; Rothenberg and Fernandez-Gonzalez, 2019). The mammalian secondary palate arises from bilateral palatal shelf anlagen of the maxillary processes that undergo outgrowth, elevation above the tongue, horizontal growth, and ultimately fusion with one another to form the intact roof of the mouth, separating the oral and nasal cavities (Ferguson, 1988; Lan et al., 2015). The medial edge epithelium (MEE) of the palatal shelves that will undergo fusion comprises an outer layer of Keratin 6a (Krt6a)-expressing squamous periderm cells that protect the palatal shelves against premature fusion and an inner layer of basal, cuboidal cells expressing the transcription factor ΔNp63 (Richardson et al., 2014); (Fitchett and Hay, 1989; Hammond et al., 2019). After the palatal shelves meet (E14.5 in mice), the opposing layers of MEE form an intervening midline epithelial seam (MES) that must be removed in order to achieve confluence of the underlying mesenchyme. Failure to complete secondary palate fusion can result in an overt cleft palate or in a submucous cleft palate, in which the oral mucosa is superficially intact but fails to form a continuous underlying structure. Cleft palate is amongst the most common human congenital anomalies, occurring in around 1/1500 births (Dixon et al., 2011). Nasolabial or median palatal epithelial cysts can also result from the enclavement of epithelium during failed embryonic palate fusion (Cinberg and Solomon, 1979; Kuriloff, 1987; Queiroz et al., 2011)

The mechanisms by which this tissue fusion occurs have been the subject of substantial investigation but remain mysterious at the cellular level. The current predominant model holds that the removal of the MES and completion of secondary palate fusion is driven mostly by epithelial apoptosis. (Carette and Ferguson, 1992; Clarke, 1990; Cuervo and Covarrubias, 2004; Cuervo et al., 2002; Hammond et al., 2019; Huang et al., 2011; Jin and Ding, 2006; Kim et al., 2015; Richardson et al., 2017; Shapiro and Sweney, 1969; Xu et al., 2008). Apoptosis plays crucial roles in many aspects of organogenesis, and its regulation can be categorized, according to apoptotic stimuli, into intrinsic and extrinsic pathways (Green, 1998). The intrinsic pathway is activated from within the cell in response to diverse signals and involves the activation of pro-apoptotic BCL-2 family proteins BAX, BAK, and BOK which regulate the mitochondrial release of cytochrome *c* and formation of a protein complex called the apoptosome, which includes the protein APAF-1. The apoptosome initiates a caspase cascade that culminates in the proteolytic cleavage and activation of the effector Caspases 3, 6, and 7 (Fuchs and Steller, 2011). The extrinsic pathway is initiated by the activation of death receptors, which recruit multiple adaptors ultimately converging on Caspase 8 to activate the same effector caspases. Common to both apoptotic mechanisms, effector caspases orchestrate destruction of the cell by cleavage of vital proteins (Danial and Korsmeyer, 2004; Nagata, 1997). Other forms of cell death include necrosis and necroptosis; though these do not involve the same cleaved caspase cascade, they ultimately converge on some similar outcomes, such as DNA fragmentation (Vanden Berghe et al., 2010).

Numerous reports observed significant apoptosis in the MES during fusion stages, and apoptosis correlates with the capacity for secondary palate fusion in some mutants, such as those with perturbed TGFβ3 signaling (AlMegbel and Shuler, 2020; Cuervo and Covarrubias, 2004; Huang et al., 2011; Iwata et al., 2011; Ke et al., 2019; Lan et al., 2015; Martínez-Álvarez et al., 2000; Shapiro and Sweney, 1969). Consistent with a requirement of apoptosis in secondary palate fusion, pan-Caspase inhibitor treatment resulted in reduced palate fusion in culture in several studies (Cuervo and Covarrubias, 2004; Cuervo et al., 2002; Huang et al., 2011), and genetic disruption of apoptosis in *Bok^-/-^;Bax^-/-^;Bak^-/-^* mutants results in a cleft palate phenotype, though loss of apoptosis in this model was not restricted to the epithelium, and these mutants also exhibited a cleft face phenotype that developmentally precedes the fusion step of secondary palatogenesis (Ke et al., 2018). On the other hand, conflicting results have been reported; pan-Caspase inhibitor treatment did not disrupt palate fusion in some studies (Takahara et al., 2004), and loss of apoptotic regulator APAF1 has been reported to result in a cleft palate in some studies (Cecconi et al., 1998; Honarpour et al., 2000) but not in others (Jin and Ding, 2006), and APAF1-independent mechanisms of cell death exist (Nagasaka et al., 2010), leaving the role of the intrinsic apoptotic pathway in secondary palate fusion uncertain. Extrinsic apoptosis mediated by the Fas ligand (FasL) has also been proposed to drive epithelial cell death during secondary palate fusion, and FasL expression was lost upon genetic perturbation of TGFβ signal reception in the palatal epithelium in mouse embryos (Huang et al., 2011; Xu et al., 2020). In addition, extensive cell extrusion has been observed during removal of the intervening MES, though whether extruding cells were all apoptotic or also included live cells was not clear (Kim et al., 2015; Schüpbach and Schroeder, 1983; Schüpbach et al., 1983).

Epithelial migration has also been proposed to contribute to the elimination of the MES cells (Carette and Ferguson, 1992; Jin and Ding, 2006; Kim et al., 2015; Logan and Benson, 2020; Richardson et al., 2017). It has been reported that TGFβ3 signaling downregulates ΔNp63 to cause basal cell cycle arrest and enable periderm cell migration to the oral and nasal aspects of the palatal shelves, which is thought to reveal the underlying basal epithelium and facilitate cell death (Cuervo and Covarrubias, 2004; Hammond et al., 2019; Richardson et al., 2017). However, epithelial migration has not been characterized and its relative functional significance remains unproven. Using live imaging, we previously proposed a mechanism of convergent displacement to explain observed movement of epithelial cells to the oral surface, but our understanding of this cell migratory mechanism was technically limited by the achievable depth of imaging with that approach (Kim et al., 2015).

Here we present new approaches for the study of secondary palate fusion, which we use to functionally interrogate the cellular behaviors that drive this process. Consistent with previous reports, we find that apoptosis is abundant within the MES before and during its removal, supporting its involvement with normal secondary palate development; however, completely blocking cell death in the epithelium only resulted in a slight delay of MES removal. Instead, through a combination of novel static- and live-imaging approaches, we uncover a surprising progression of collective epithelial cell migratory patterns. Small gaps in the MES consolidate into an interconnected network of epithelial trails that connect to the oral and nasal surfaces, and epithelial islands that undergo apoptosis, or migrate through the mesenchyme. Whereas adherens junctions couple epithelial cells within trails during migration, filamentous actin is anisotropically enriched at the edges of trails and contractility promotes their collective peristaltic movement. Actomyosin contractility is critical for this unique form of epithelial migration, and its disruption results in the dissolution of epithelial collectives and failure to complete secondary palatal shelf fusion normally. These results provide new insight into the cellular mechanisms driving secondary palate fusion and indicate that redundant cellular processes mediate this crucial morphogenetic event.

## RESULTS

### A new imaging approach shows that apoptosis is not required for MES removal during palate fusion

To characterize cell behaviors during secondary palate fusion, we first aimed to better visualize the MES. As traditionally-employed coronal sections provide a view of the MES at one anteroposterior position, we established a sagittal thick-sectioning technique to enable visualization of the MES in its entirety (Fig. S1A-D). Static imaging of MES thick sections immunostained for E-cadherin at progressive stages of MES removal revealed a surprising pattern of MES clearance (Fig. 1A-C). Small epithelial holes appeared just before E14.75, and widened by E15.0 to give the appearance as a web-like network of trails connecting to the oral and nasal surface epithelium (Fig. 1A-C). Small epithelial islands which appeared to have separated from the trails, were particularly apparent at E15.5 (Fig. 1C). At each of these stages, cleaved Caspase-3 immunostaining revealed that between ten and forty percent of MES cells undergo apoptosis, consistent with the prevailing understanding that apoptotic death is the ultimate fate of a substantial proportion of MES cells during normal secondary palate fusion (Cuervo and Covarrubias, 2004; Cuervo et al., 2002). Further, we found that a greater proportion of MES cells in the posterior palate undergo apoptosis compared with more anterior positions (Fig. 1D-F). This new imaging perspective inspires a new investigation into the patterns of MES removal during secondary palate development.

**Figure 1.**
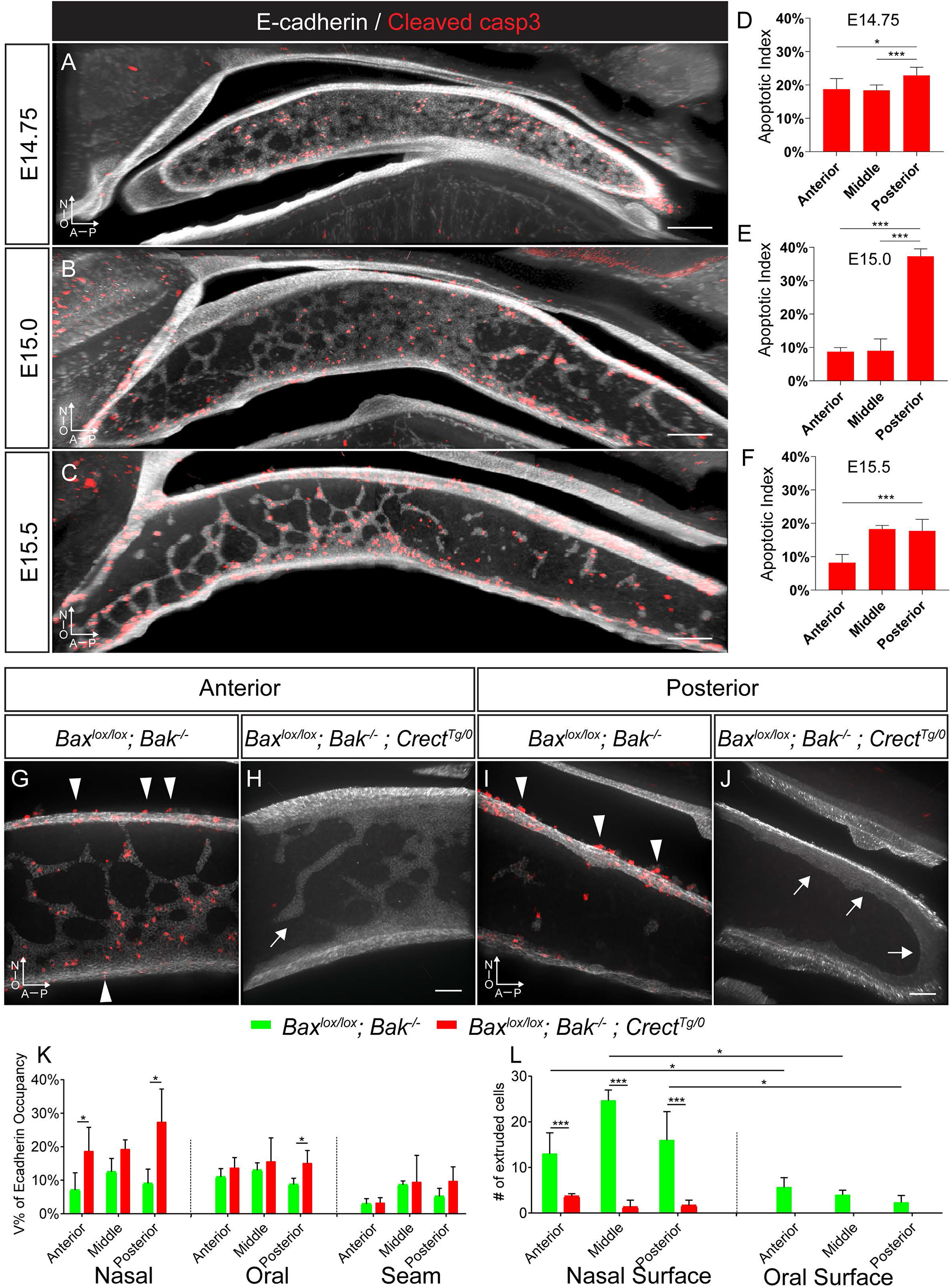
Apoptosis is abundant in the MES but its loss does not prevent MES removal. (A-C) 3D-rendered views of sagittal sections of wild-type mouse secondary palate immunostained for E-cadherin (white) and Cleaved Caspase-3 (red). (A) At E14.75, the E-cadherin-expressing MES is organized in a continuous sheet. (B) At E15.0, gaps in the MES are larger and create a network of epithelial trails connecting to the oral and nasal epithelial surfaces (C) By E15.5, the remaining MES is organized into a series of interconnecting trails; at this stage epithelial islands have sometimes broken away from epithelial trails. At each of the above stages, Cleaved Caspase-3 immunostaining reveals the abundance of MES apoptosis. A, anterior; P, posterior; N, nasal surface; O, oral surface. Scale bar, 200μm. (D-F) Quantification of apoptosis at each stage, and across distinct anteroposterior regions during fusion. Histograms show the apoptotic index, which is the ratio of the number of apoptotic MES cells to the total number of MES cells in entire secondary palate at each stage. Column height represents the means of the ratio for the number quantified per stage. Error bars represent S.E.M, *, P<0.03; ***, P<0.001. Number of embryos analyzed is presented in Table S1. (G-J) 3D-rendered images of (G,H) E15.5 mid-anterior and (I,J) posteror regions of (G,I) *Bax^lox/lox^; Bak^-/-^* and (H,J) *Bax^lox/lox^; Bak^-/-^; Crect^Tg/0^* sagittal thick sections immunostained for E-cadherin (white) and Cleaved Caspase-3 (red). Arrowheads indicate apoptotic cell extrusion events in *Bax^lox/lox^; Bak^-/-^* embryos, which are lost in *Bax^lox/lox^; Bak^-/-^; Crect^Tg/0^* embryos, which also exhibit thickened MES trails (arrows). Scale bar, 50μm. (K-L) Quantification of E-cadherin occupancy (K) and cell extrusion (L) in indicated palatal regions. Column height represents the mean for a given genotype, error bars represent S.E.M., *, P<0.03; ***, P<0.001. Number of embryos analyzed is presented in Table S1.

We employed this imaging approach to study the function of apoptosis within the MES without confounding effects of earlier cranial and neural malformations, utilizing *Crect* Cre craniofacial ectoderm driver to disrupt the *Bax* gene specifically in the MES of *Bak* mutant mice (Reid et al., 2011). Compared with *Bax^lox/lox^; Bak^-/-^* embryos that exhibited extensive apoptotic cell death, we found that cell death was completely lost in the MES of *Bax^lox/lox^; Bak^-/-^; Crect^Tg/0^* E15.5 mutant embryos, assayed by both Cleaved Caspase-3 immunostaining and TUNEL analysis (Fig. 1G-J; Fig. S1E-F). Quantifying the volume of E-cadherin-expressing MES cells at E15.5 revealed that whereas loss cell death did not prevent secondary palate fusion, it did appear to change the pattern and timing of MES cell removal. This effect was most substantial within the posterior secondary palate, where the MES was retained in the oral, posterior, and nasal surfaces of the posterior palate (Fig. 1I-K), consistent with the greater amount of MES apoptosis in this region. Cell extrusion was abundant on the oral and nasal surfaces of the secondary palates of controls (Fig. 1G, I, L), but nearly completely lost in *Bax^lox/lox^; Bak^-/-^; Crect^Tg/0^* E15.5 mutant embryos (Fig. 1H, J, L), indicating that the apoptotic form of cell extrusion predominates, but that its loss does not result in a failure of MES removal. Together, these results indicate that the vast majority of cell death within the MES is attributable to the intrinsic apoptotic pathway, that cell extrusion is largely apoptotic, and that upon loss of both of these cellular mechanisms, secondary palate fusion and MES clearance completed successfully. Therefore, though these mechanisms may normally contribute to MES removal, their loss can be overcome by other cellular mechanisms.

Epithelial to mesenchymal transition (EMT) has also been proposed to contribute to MES removal, but genetic lineage tracing did not reveal a contribution of MES cells to the underlying mesenchyme (Fitchett and Hay, 1989; Gritli-Linde, 2007). Nevertheless, we wondered if loss of apoptosis might result in a compensatory increase in EMT to clear MES cells. We performed genetic lineage tracing of the epithelium in *Bax^lox/lox^; Bak^-/-^; Crect^Tg/0^*; *R26^mTmG/+^* and control *Crect^Tg/0^*; *R26^mTmG/+^* embryos at E15.5. The *R26^mTmG^* reporter enables the expression of membrane-tethered GFP following Cre-mediated recombination (Muzumdar et al., 2007). We did not observe any GFP reporter positive, E-cadherin negative cells throughout most of the palate, in the presence or absence of apoptosis, indicating that compensatory EMT is not at play (Fig. S2A, C). In contrast, we did detect GFP-expressing cells in the mesenchyme of the most posterior aspect of the secondary palatal shelves in both genotypes (Fig. S2B, D). However, *in situ* hybridization by RNAscope one day earlier (E14.0) revealed an abundance of *Cre* transcript expression, indicating that the *Crect* lineage in the posterior palate mesenchyme was attributable to unexpected *Crect* activity (Fig. S2E, F). These data suggest no compensatory EMT in most of the secondary palate, but our ability to detect EMT in the far posterior palate was obfuscated by the mesenchymal activity of the *Crect* allele in this region. This activity will be an important consideration in future studies of secondary palate development that make use of this nonetheless valuable mouse line (Reid et al., 2011).

### A unique form of collective epithelial migration drives secondary palate fusion

These patterns of epithelial removal suggested that collective cell migration may play a central role. We previously published a system for *ex-vivo* live imaging of secondary palate fusion, which employed laser scanning or spinning disk confocal microscopy to directly image whole secondary palatal shelf explant cultures from the oral side of the MES in fluorescent reporter mouse embryos (Kim et al., 2015; Kim et al., 2017). This approach was limited by the achievable depth of focus in live tissues, as only the most superficial cells at the oral aspect of the MES (known as epithelial triangles) could be visualized. We therefore applied our sagittal thick section imaging technique to spinning disk confocal imaging in order to perform *ex-vivo* live imaging of MES removal in the secondary palate in *Crect^Tg/0^; R26^mTmG/+^* embryos. We initiated imaging at multiple time points between E14.75 and E15.5 in order to better understand epithelial cell movements during MES removal. First, by live imaging the mid-anterior MES for 20 hours starting at E14.75, we observed initial small breaks in the epithelium that enlarged as surrounding epithelial cells underwent rearrangements, breaking and re-establishing epithelial junctions until coalescing ultimately into a network of trails similar to what we described from our static imaging at E15.5 (Movie 1). We noticed extensive MES cell blebbing over this time-course, consistent with the occurrence of apoptosis (Fig. 2A; Movie 1). Live imaging starting at E15.5 revealed that epithelial trails streamed continuously toward the surface epithelium (Fig. 2B; Movie 2). To determine whether epithelial cells moved as a collective, we utilized the nuclear *R26^nTnG^* Cre reporter to track individual nuclei over the course of eight hours. Tracking individual cells revealed that though some rearrangements occurred within epithelial streams, the nuclei moved collectively from positions deep within the seam to the oral or nasal epithelium (Fig. 2E, Movie 3). During their migration, epithelial trails often broke into smaller cell collectives, ostensibly forming the previously-described epithelial islands (Fig. 2B). When epithelial islands were close to the surface epithelium, they ultimately contacted and coalesced with the oral or nasal surface epithelium (Fig. 2C, Movie 4). However, epithelial islands that were deep within the secondary palatal shelves exhibited apoptotic bodies and membrane blebbing as they progressively disappeared through apparent cell death over the course of imaging (Fig. 2D; Movie 5).

**Figure 2.**
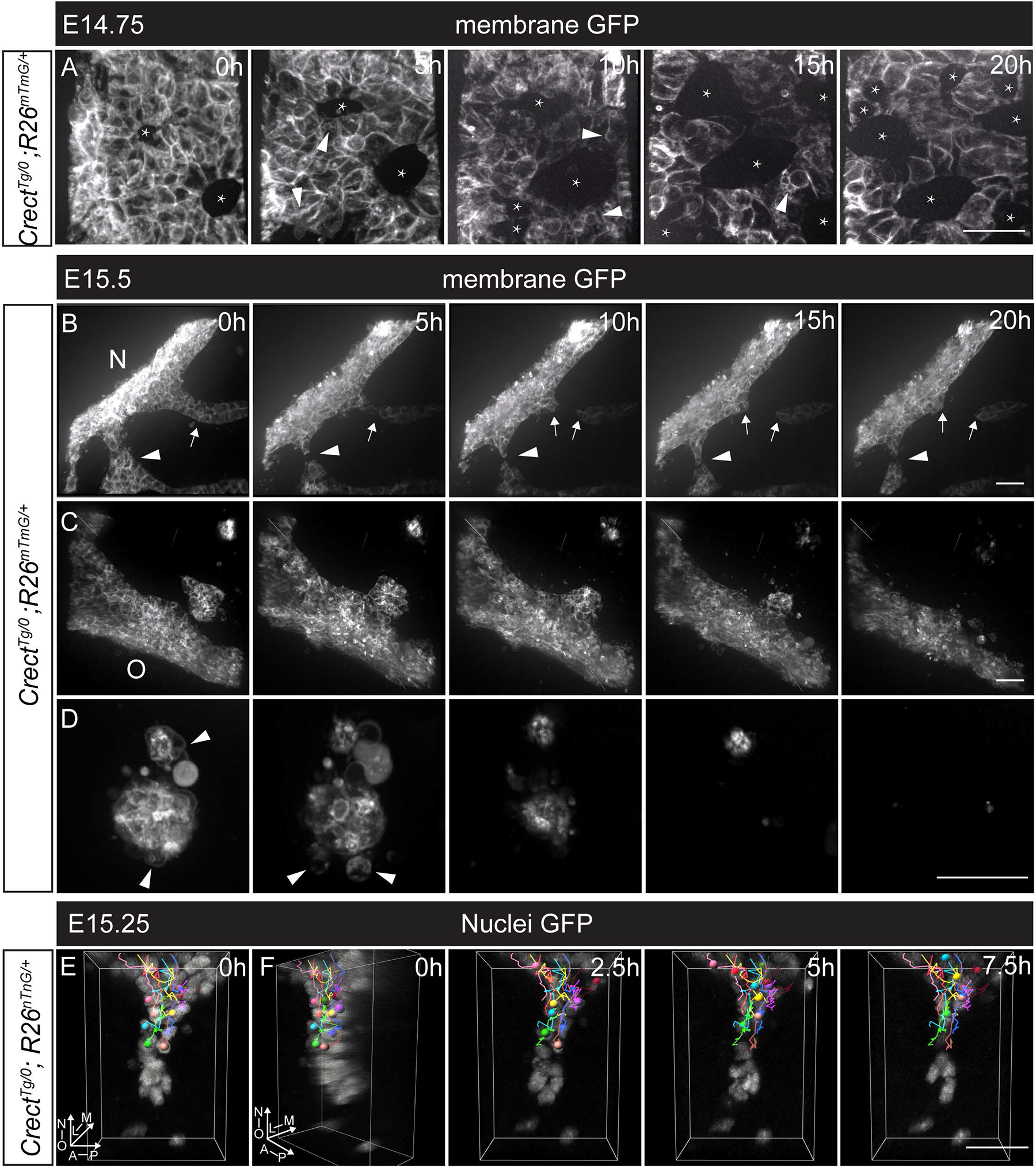
A novel live imaging approach reveals patterns of collective epithelial migration in MES removal. (A-D) Confocal live imaging of EGFP in *Crect^Tg/0^; R26^mTmG/+^* E14.75 and E15.5 embryos in various regions of the secondary palate of reveals cell behaviors during MES removal. (A) Starting from E14.75, the MES “sheet” exhibited small breaks which enlarged into holes (indicated by asterisks), and arrowheads point to membrane blebbing. Scale bar, 25 (See also Movie 1) (B) Live imaging beginning at E15.5 revealed cell behaviors associated with the clearance of MES trails. An example of an MES cell trail moving toward the nasal surface as an epithelial collective (arrowheads); trails often break to form smaller trails or epithelial islands (arrows).(See also Movie 2) (C) An example of an epithelial island close to the oral surface as it coalesces into the oral surface epithelium. (See also Movie 4) (D) MES cells in an island far from the nasal or oral surface undergoes apoptosis, arrowheads point to membrane blebbing, as the island progressively shrinks and disappears. Scale bars, 25μm. (E-F) Confocal live imaging and cell tracking of cells in an EGFP-labeled trail in a E15.25 *Crect^Tg/0^; R26^nTnG/+^* embryo. (See also Movie 5) (F) image in E turned 45° in the oral-nasal axis. Spots were created and manually traced using Imaris and spots and tracks were labeled by tracking ID color. MES trail cells, indicated by colored spots, were tracked during a 7.5 hours imaging period. Lines indicate the movement of each cell. Scale bar, 25 μm. h, hour; N, nasal surface; O, oral surface; L, lateral; M, medial; A, anterior; P, posterior.

Based on the fact that the MES was successfully removed in the absence of cell death, but that patterns of epithelial removal were altered, we performed live imaging of *Bax^lox/lox;^ Bak^-/-^; Crect^Tg/0^; R26^mTmG/+^* mutant embryos to observe whether the cell behaviors underlying MES removal were altered upon loss of apoptosis. Epithelial breakage, and collective migration of epithelial trails in mutants appeared overtly similar to control embryos, though we observed reclosure of some epithelial holes that we did not see in wild-type embryos (Fig. 3A,B; Movie 6,7). Epithelial islands in mutants underwent cell blebbing similar to control but failed to shrink or disappear (Fig. 3C; Movie 8; Fig. 2D; Movie 4). These persistent epithelial islands together with occasional recovery of epithelial breakage may account for the mildly altered epithelial removal seen in *Bax^lox/lox;^ Bak^-/-^; Crect^Tg/0^* mutants. Overall, our results indicate that in the absence of apoptosis, MES cells retain collective epithelial migration behaviors that compensate for loss of apoptotic contributions to MES removal.

**Figure 3.**
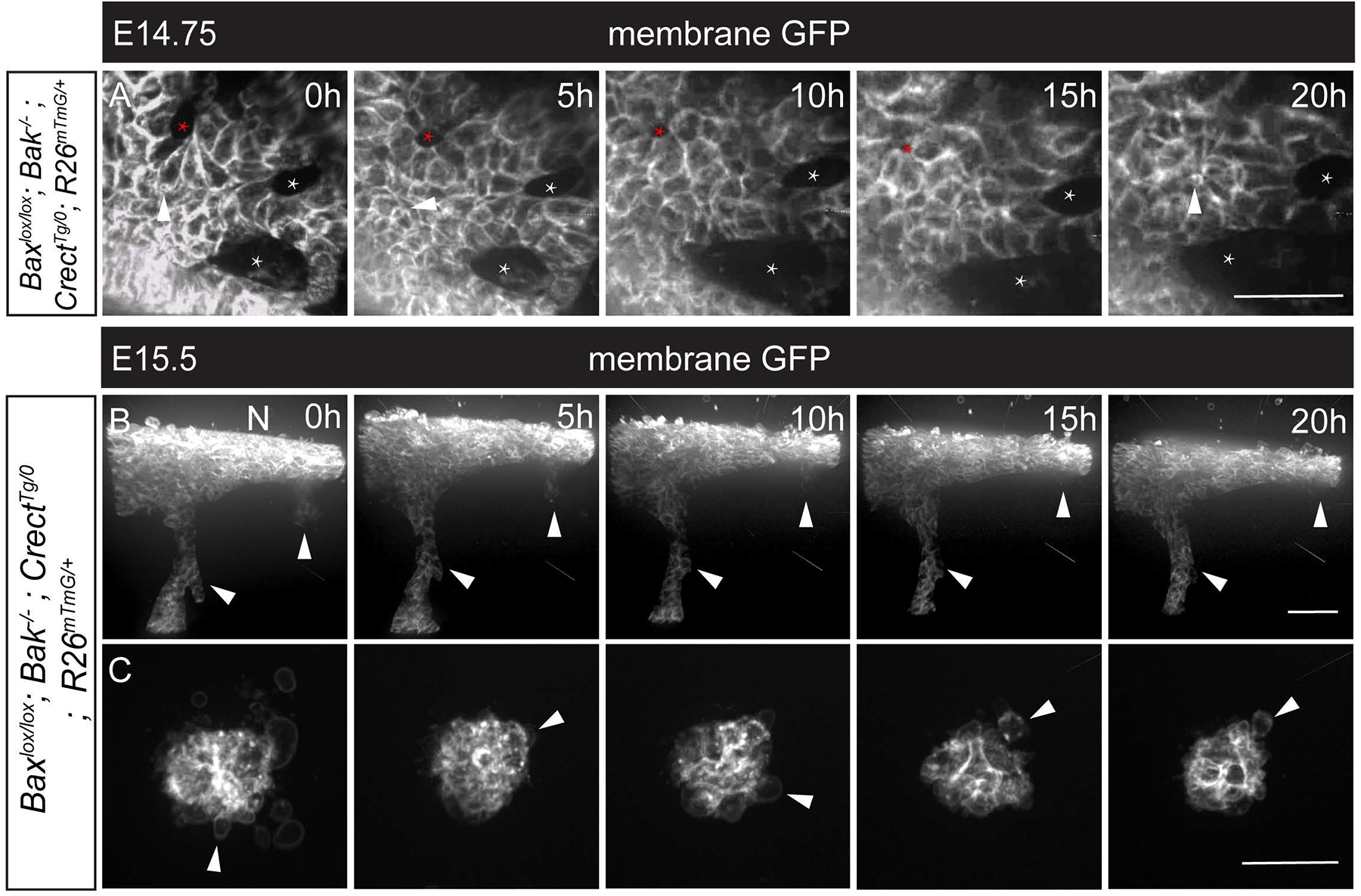
Loss of BAX and BAK results in changes in MES cell removal, but retention of collective epithelial cell migration. (A-C) Confocal live sagittal section imaging of *Bax^lox/lox^; Bak^-/-^; Crect^Tg/0^; R26^mTmG/+^* embryos at E14.75 and E15.5 reveals MES cell behaviors upon loss of apoptosis. (A) Over a 20-hour imaging period starting at E14.75, dynamic cell rearrangements within the MES epithelium result in the expansion of some holes (white asterisk), and the closure of another (red asterisk); arrowheads point to membrane blebbing. (See also Movie 6) (B) MES cells migrate as collective trails resulting in MES cells being incorporated into the nasal surface epithelium (arrowheads, Gamma 1.5, See also Methods). (See also Movie 7) (C) Islands found deep within the palatal shelves did not disappear over a 20-hour imaging period, but did exhibit membrane blebbing (arrowheads) (See also Movie 8) N, Nasal surface. Scale bar, 25 μm.

### Krt6a/p63 expressing epithelial cells undergo collective migration

Based on a previous report indicating that periderm cells undergo migration during palatal fusion (Richardson et al., 2017), and to discern between cell behaviors of peridermal vs. basal MES cells, we generated a *Krt6a iCre* knock-in mouse line for tracking the migration of peridermal cells. Crossing *Krt6a^iCre^* mice with the *R26^nTnG^* nuclear reporter allele confirmed that recombination was restricted to the periderm and was entirely non-overlapping with the ΔNp63-expressing basal epithelial cells prior to palatal shelf contact at E14.0 (Fig. 4A). Sagittal sections of the MES undergoing fusion at E15.0 showed that epithelial cells that are migrating to the nasal surface (Fig. 4B), within the medial aspect of the secondary palate (Fig. 4C), or migrating to the oral surface (Fig. 4D) all exhibited GFP expression reflecting *Krt6a* lineage identity. *Krt6a^iCre^; R26^nTnG^* periderm lineage positive cells that did not express p63 mostly resided within the “epithelial triangles” of the nasal surface. Surprisingly, in migratory epithelial trails, we observed co-expression of the *Krt6a^iCre^; R26^nTnG^* periderm lineage marker together with p63. As our finding that p63 expression was retained within MES cells during fusion differs from what has been previously published (Richardson et al., 2014; Richardson et al., 2017), we compared staining using an antibody recognizing pan-p63 with staining using an antibody specifically recognizing the ΔNp63 isoform. These antibodies exhibited identical staining patterns suggesting that this discrepancy from previously published work does not reflect a difference in isoform expression (Fig. 4B-D). The abundance of Krt6a lineage and p63 double positive cells therefore provides evidence that p63-expressing basal epithelial cells undergo migration to the oral and nasal surface concomitant with the basal epithelial initiation of expression of periderm marker Krt6a.

**Figure 4.**
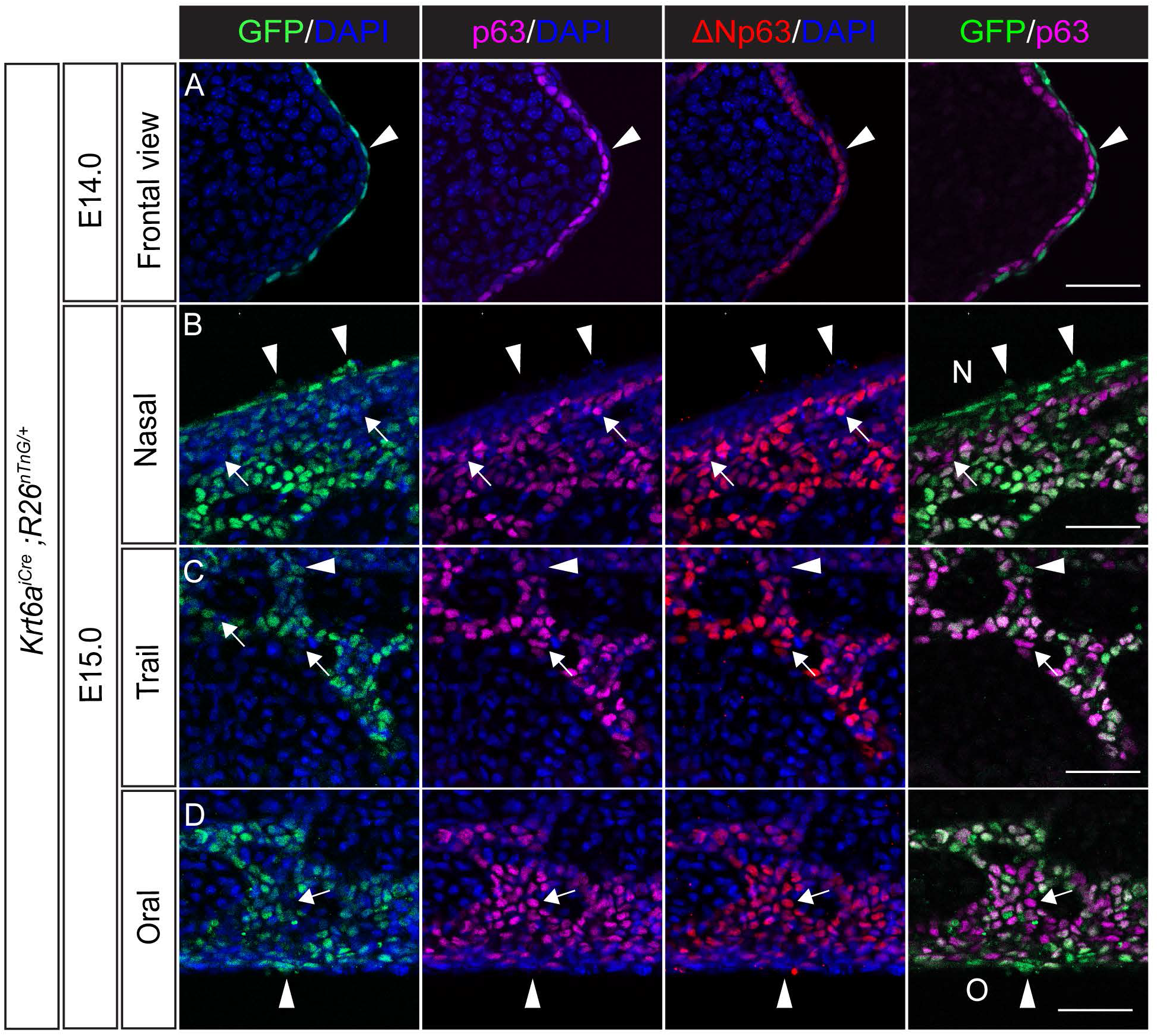
Migrating MES cells increase expression of periderm marker Krt6a while retaining basal marker p63. (A) Immunostaining of coronal cryosections of E14.0 *Krt6a^iCre^; R26^nTnG^* embryos for GFP (green), pan-p63 (magenta) and the ΔNp63 isoform (red) confirms Cre recombination activity specifically within the periderm prior to palatal shelf contact. (B-D) Optical slices of vibratome thick sagittal sections of E15.0 *Krt6a^iCre^; R26^nTnG^* embryos reveals co-expression of the Krt6a lineage (GFP) with the basal markers p63 and ΔNp63 within most cells of the migratory MES trails. p63 expression was lost only from cells in “epithelial triangle” regions close to the nasal (B) and oral (D) surfaces. Arrows point to GFP^-^p63^+^ cells, and arrowheads indicate GFP^+^p63^-^ cells. N, nasal surface; O, oral surface. Scale bars, 25μm.

### Collective epithelial migration of the MES is driven by actomyosin contractility

Cell migration requires actomyosin contractility generated by non-muscle myosin, and we previously discovered that loss of non-muscle myosin isoforms NMIIA and NMIIB, encoded by the *Myh9* and *Myh10* genes, respectively, results in a cleft secondary palate (Kim et al., 2015; Vicente-Manzanares et al., 2009). We examined loss of NMII function using sagittal sectioning and found that *Myh9^lox/lox^; Myh10^lox/+^; Crect^Tg/0^* compound mutants, which lacked NMIIA and had reduced NMIIB specifically in the MES, lost epithelial trail and island organization and instead exhibited a diffuse and dispersed epithelium at E15.5 (Fig. 5A-E). In addition to inappropriate retention of dispersed epithelium, *Myh9^lox/lox^; Myh10^lox/+^; Crect^Tg/0^* mutant embryos also exhibited a decrease in the absolute number of apoptotic cells (Fig. 5F), and a dramatic decrease in apoptotic cells relative to the total amount of retained epithelium (Fig. 5G), suggesting that actomyosin contractility might be required to stimulate MES cell death. Analysis of individual *Myh9^lox/lox^; Crect^Tg/0^* mutant embryos revealed a failure of MES removal that was similar to compound *Myh9^lox/lox^; Myh10^lox/+^; Crect^Tg/0^* mutant embryos at E15.5, whereas *Myh9^lox/+^; Myh10^lox/lox^; Crect^Tg/0^* compound mutants looked similar to controls (Fig. S3), suggesting that NMIIA is the more crucial regulator. Additionally, at E17.5, an abundance of epithelial inclusions persisted in the secondary palate of *Myh9^lox/lox^; Crect^Tg/0^*; *R26^mTmG/+^* mutants (Fig. S4), further supporting the importance of the NMIIA isoform in secondary palate fusion.

**Figure 5.**
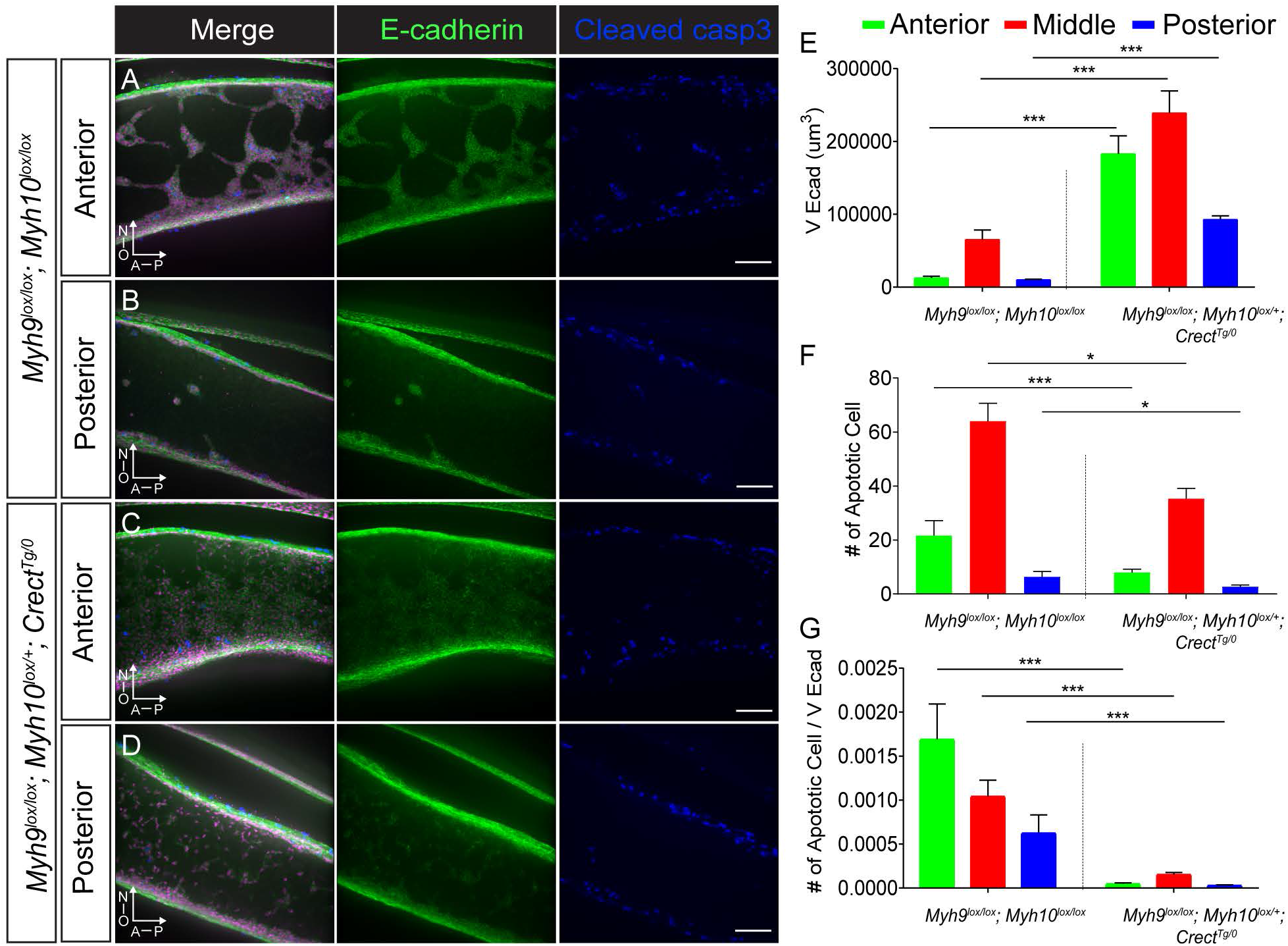
Epithelial-specific loss of NMII results in failure to clear the MES. (A-D) *Myh9^lox/lox^; Myh10^lox/lox^* control and *Myh9^lox/lox^; Myh10^lox/+^; Crect^Tg/0^* mutant sagittal sections of secondary palates immunostained for E-cadherin (green), p63 (magenta), and cleaved caspase-3 (blue) shown as 3D-rendered images. (A,B) In control, E-cadherin labeled MES cells are organized into collective trails and islands as they are removed from the palatal shelves (C,D) In *Myh9^lox/lox^; Myh10^lox/+^; Crect^Tg/0^* mutant embryos, an abundance of E-cadherin labeled MES cells remain in the palate, and epithelial cells appeared more dispersed. Scale bar, 50 μm. A, anterior; P, posterior; N, nasal; O, Oral (E) Quantification of E-cadherin expression volume (µm^3^) as a measure of MES removal in the anterior, middle, and posterior palate of *Myh9^lox/lox^; Myh10^lox/lox^* control and *Myh9^lox/lox^; Myh10^lox/+^; Crect^Tg/0^* mutant secondary palates (F) Number of apoptotic cells in the anterior, middle, and posterior palate of *Myh9^lox/lox^; Myh10^lox/lox^* control and *Myh9^lox/lox^; Myh10^lox/+^; Crect^Tg/0^* mutant secondary palates. (G) Apoptotic cell number normalized by total E-cadherin volume in the anterior, middle, and posterior palate of *Myh9^lox/lox^; Myh10^lox/+^; Crect^Tg/0^* and control embryos. Error bars represent S.E.M., *, P<0.03; ***, P<0.001. Number of embryos analyzed is presented in Table S1.

To determine how loss of NMII activity affected epithelial cell behaviors underlying secondary palate fusion, we performed live imaging of *Myh9^lox/lox^; Crect^Tg/0^*; *R26^mTmG/+^* mutant embryos and compared them with controls that still preserved NMII function (Fig. 6A, B). Whereas controls exhibited streaming collective epithelial migration (Fig. 6A; Movie 9), complete loss of NMIIA in *Myh9^lox/lox^; Crect^Tg/0^*; *R26^mTmG/+^* mutants resulted in complete loss of epithelial trail organization and GFP positive MES-lineage cells that traveled in a disorderly fashion, had more protrusive shapes, and were ultimately not cleared from the palatal shelf mesenchyme (Fig. 6B; Movie 10). At E15.5, highly-ordered E-cadherin junctions are present between epithelial cells but not at the epithelial-mesenchymal cell interface in control MES cell trails (Fig. 6C; S3A, B). In contrast, GFP positive MES-lineage cells lacking NMIIA in mutants exhibited a dramatic loss of junctional E-cadherin, which also appeared reduced in abundance, possibly due to destabilization by loss from junctions (Fig. 6D; S3C-F). Interestingly, loss of epithelial architecture resulted in a highly protrusive, almost mesenchymal appearance of GFP positive MES lineage cells in *Myh9^lox/lox^; Crect^Tg/0^; R26^mTmG/+^* mutant embryos, suggesting that NMIIA is required to maintain proper polarization of epithelial collectives (Fig. 6D; S3C-F)(Pandya et al., 2017). At the later timepoint of E17.5, GFP positive MES lineage cells still retained E-cadherin, indicating that in the absence of NMIIA, the MES loses epithelial organization and junctions, exhibits reduced E-cadherin, and drastically changes its migratory cell behaviors; yet these cells do not stably contribute to the palatal shelf mesenchyme by EMT in the majority of the secondary palate (Fig. S4A-D).

**Figure 6.**
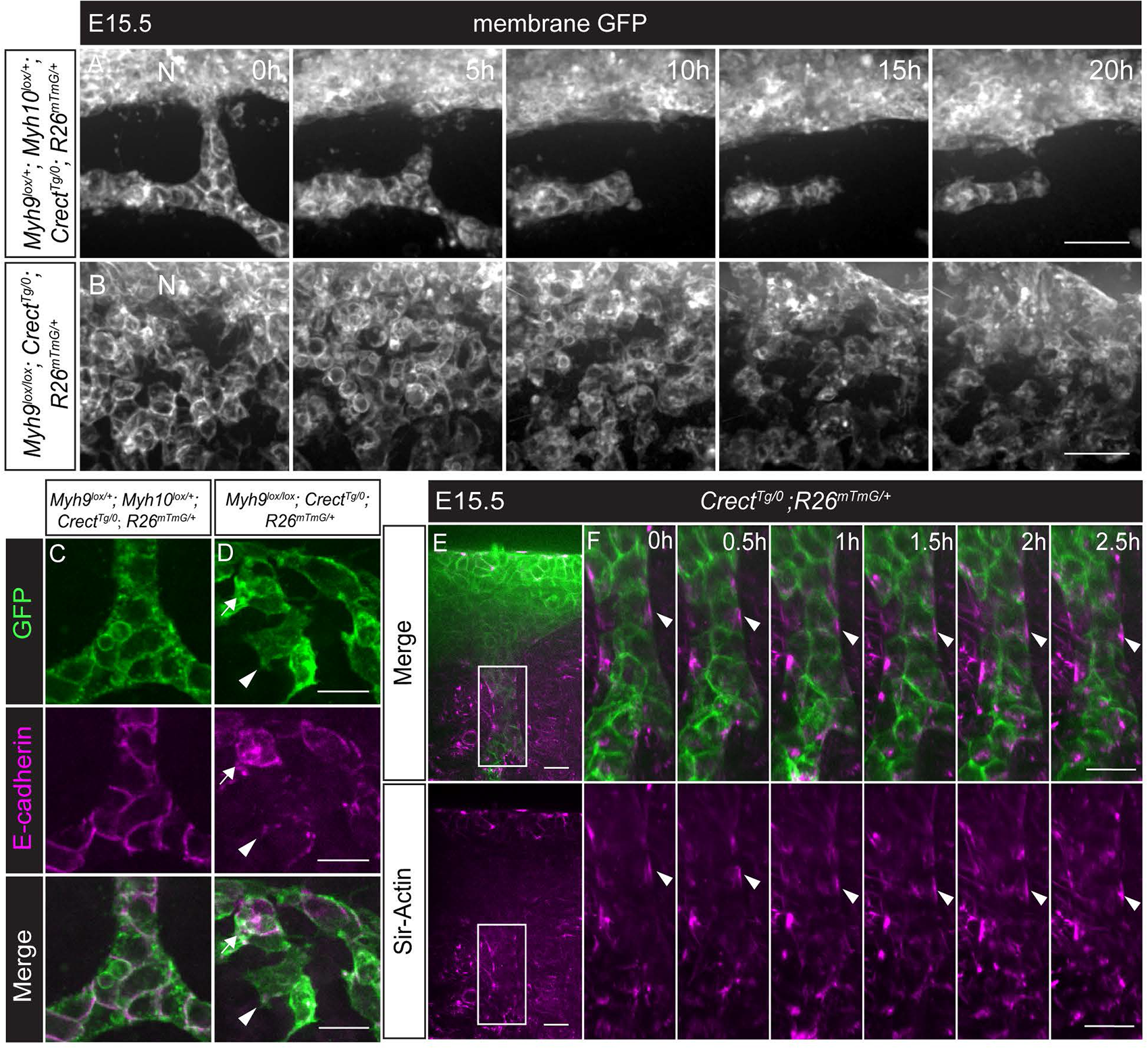
NMII is required for collective epithelial organization and peristaltic movement of MES trails. (A-B) Confocal live imaging of *Myh9^lox/+^; Myh10^lox/+^; Crect^Tg/0^; R26^mTmG/+^* control and *Myh9^lox/lox^; Crect^Tg/0^; R26^mTmG/+^* mutant sagittal sections at E15.5. (A) In controls, a trail of cells moves towards the nasal surface and breaks, resulting in island formation. (See also Movie 9) (B) In the mutant, MES cells lose trail and island organization and cells take on a more protrusive shape and move in an undirected manner. (See also Movie 10) N, nasal surface. Scale bar, 20μm. (C-D) 3D rendering of immunostained MES trails at high magnification in E15.5 control (C) and *Myh9^lox/lox^; Crect^Tg/0^; R26^mTmG/+^* mutant (D) embryos reveals a loss of junctional E-cadherin and an apparent reduction in E-cadherin levels upon loss of NMIIA. Scale bar, 5μm. White arrow points to a cell retaining high levels of non-junctional E-cadherin; white arrowhead points to a cell with loss of adherens junction and reduced E-cadherin level (magenta) (E-F) Sagittal thick sectioning confocal live imaging of a E15.5 *Crect^Tg/0^; R26^mTmG/+^* secondary palate. EGFP (green) labeling of MES cells and SiR-Actin (magenta) labeling of F-actin reveals anisotropic F-actin accumulation in epithelial trails. Pulsatile contractility of actomyosin drives the movement of the epithelial trail toward the nasal surface. (See also Movie 11) (E) Low magnification views of a single optical section. (F) 3D rendering of the boxed region in (E) at high magnification. Arrowheads follow the movement of one F-actin filament. Scale bars, 15 μm.

To better understand how NMII regulates collective epithelial migration, we performed live imaging of *Crect^Tg/0^; R26^mTmG^* embryos treated with the SiR-Actin spirochrome dye to directly observe filamentous actin over the course of MES removal. We found that epithelial trails undergoing migration to the nasal and oral epithelium exhibited anisotropic distribution of filamentous actin cables at the epithelial-mesenchymal interface (Fig. 6E-F). As epithelial trails underwent collective movement toward the nasal and oral epithelium, these actomyosin cables contracted in a pulsatile fashion and appeared to move epithelial trails in a peristaltic fashion toward the oral and nasal edges (Fig. 6E-F; Movie 11). These data reveal a unique form of collective epithelial migration that is responsible for MES clearance during secondary palate fusion.

## DISCUSSION

The cellular mechanisms driving secondary palate fusion have been investigated for more than three decades, leading to the thought that apoptosis is a crucial final step in the removal of the MES. Indeed, using our novel sagittal imaging methods, we observed an abundance of apoptosis and apoptotic cell extrusion in MES cells during their removal. Further, live imaging of epithelial islands confirms epithelial loss through cell death, consistent with cell death being the ultimate fate of a substantial proportion of MES cells, particularly in the posterior palate. Loss of intrinsic apoptotic regulators BAX and BAK resulted in a complete loss of MES cell death and prevented cell extrusion, indicating that cell death in the MES occurs through the intrinsic apoptotic pathway, and that the bulk of extruded cells are apoptotic. Surprisingly though, complete loss of cell death and cell extrusion did not dramatically disrupt MES clearance or secondary palate fusion, which, we herein describe, proceeds through a progressive series of epithelial cell movements. First, small breaks in the MES sheet enlarge through the rearrangement of surrounding epithelial cells, and epithelia coalesce into a web-like network of trails and islands. The trails migrate as epithelial collectives, where they incorporate into the oral and nasal surfaces, or are eliminated through apoptotic cell extrusion. Cell migratory behaviors are maintained in the absence of apoptosis and cell extrusion and collective epithelial cell migration overcomes the loss of apoptosis to carry out MES cell removal (Fig. 7).

**Figure 7.**
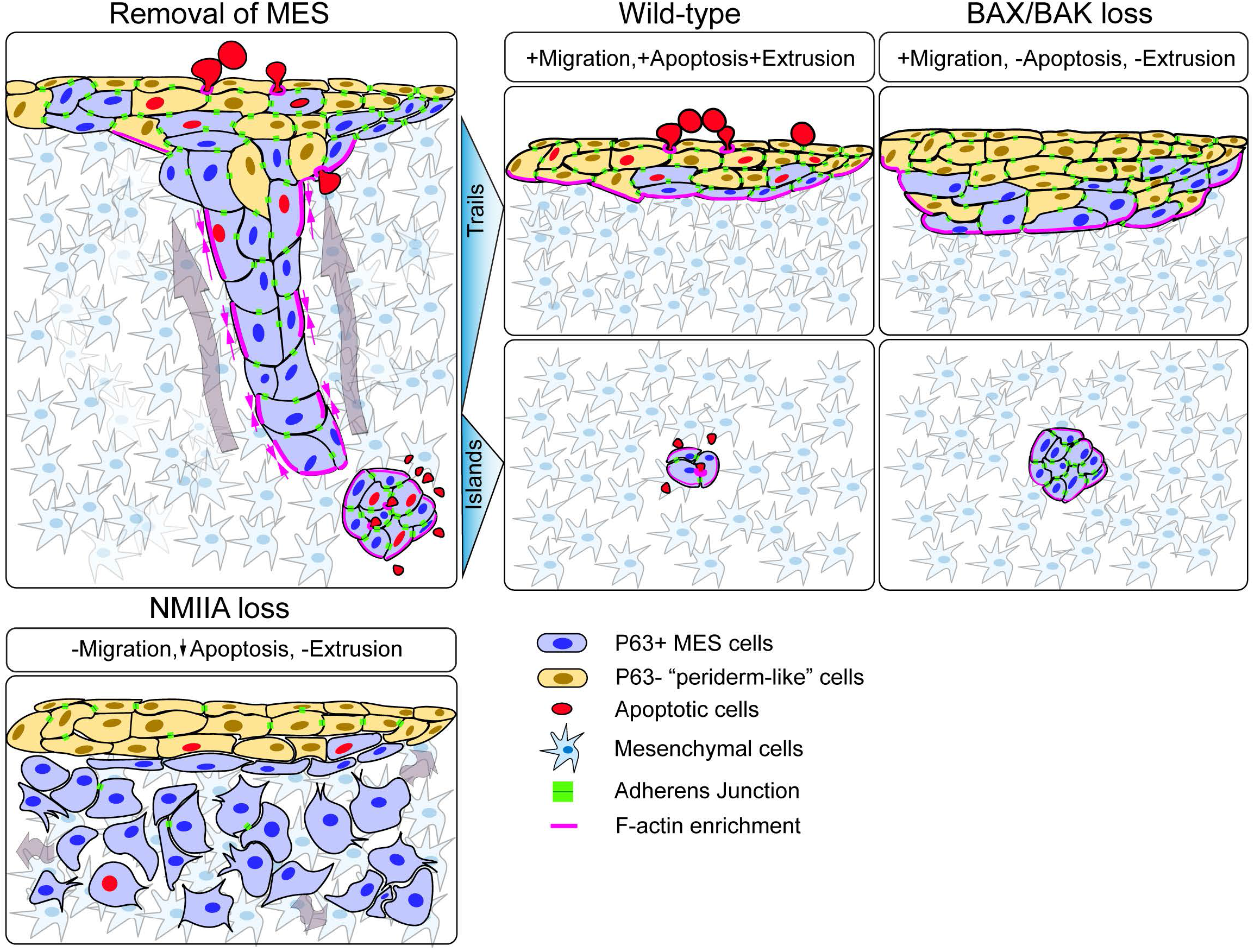
Cell behaviors driving MES removal during secondary palate fusion. Collective migration of MES cells occurs through the streaming migration of trails, which sometimes break into islands. Trails are removed through actomyosin contractility dependent peristaltic collective movement, whereas islands are removed through either apoptosis or migration. Loss of actomyosin contractility through mutation of NMIIA results in dispersal of epithelial collectives and failure to clear the MES. Loss of function of BAX and BAK result in subtle changes in MES removal and epithelial islands do not shrink, but instead migrate toward the oral and nasal surfaces; loss of apoptosis does not prevent completion of MES removal.

Previous attempts to observe cell behaviors during MES removal included static studies using *in vitro* culture of unpaired palatal shelves, which led to the conclusion that MES clearance does not depend on contact from the opposite palatal shelf (Charoenchaikorn et al., 2009; Takigawa and Shiota, 2004; Yamamoto et al., 2020). The patterns of epithelial removal that were described using these protocols, however, were quite unlike what we observed using our approaches, suggesting that the cellular behaviors underlying epithelial loss in those models may be impacted by cell culture conditions. Indeed, unpaired palatal shelf culture performed in media that included amniotic fluid did not result in loss of epithelium (Takigawa and Shiota, 2004), which is also consistent with the fact that there are many mouse mutants that exhibit cleft palate due to defects in palatal shelf outgrowth, thereby resulting in a loss of palatal shelf contact, but which retain the palatal epithelium (Bush and Jiang, 2012). Taken together, the evidence is consistent with MES cell apoptosis and migration requiring palatal shelf contact for their initiation, but the molecular cues initiating migration remain unknown (Carette and Ferguson, 1992).

It has been repeatedly demonstrated that apoptosis does not occur upon loss of TGFβ3 signaling, which converges on Smad-dependent and Smad-independent pathways to regulate palate fusion (Lane et al., 2015; Xu et al., 2008). As TGFβ signaling is absolutely required for MES clearance, but apoptosis is not, TGFβ3 must also regulate other cell behaviors in this process. Indeed, TGFβ3 signaling regulates the differentiation of the MES, including the downregulation of ΔNp63 which in turn regulates numerous cell adhesion genes (Richardson et al., 2017). Further the compound loss of TGFβ3 and p63 in *Tgfβ3^-/-^; p63^+/-^* mutant embryos resulted in rescue of secondary palate fusion, providing functional support for this epistatic relationship (Richardson et al., 2017). We were therefore surprised to observe an abundance of cells co-expressing ΔNp63 and the periderm marker, Krt6a in migrating epithelial trails. However, the increase in expression of Krt6a that we observed within basal epithelial cells fits well with previous studies that demonstrate that, like Krt6a, Krt17 expression is also increased within the basal epithelium of the MES during fusion (Jin et al., 2014; Vaziri Sani et al., 2005). We therefore hypothesize that these cells may represent an intermediate differentiation state as cells migrate in streams toward the oral and nasal surfaces. Richardson et al. also used a transgenic mKrt17-GFP reporter to demonstrate that the Krt17-expressing periderm cell layer underwent movement during palate fusion (Richardson et al., 2017). Our results, which suggest that basal epithelial cells expressing ΔNp63 also take on periderm-like gene expression during fusion, is consistent with this observation and suggest that MES migration is concomitant to a change in basal epithelial differentiation state. It is interesting to note that Krt6a expression is also upregulated at other sites of epithelial fusion, such as the eyelids and embryonic wounds, and its loss affected migration of keratinocytes during wound healing (Mazzalupo and Coulombe, 2001; Wojcik et al., 2000). It remains to be seen whether the change in expression of peridermal keratins such as Krt6a is involved in differentiation to a migratory cell phenotype or if it is reflective of transitioning from a basal cell type to a surface cell type that must provide epithelial barrier integrity. In the future, deep characterization of spatial changes in gene expression during palatal fusion will help to illuminate how differentiation is coupled with MES migration.

Several previous studies provided evidence that cell migration occurs during secondary palate fusion, but the nature of the migration and whether it is required for fusion was not clear (Carette and Ferguson, 1992; Kim et al., 2015; Richardson et al., 2017). We demonstrate that MES cells do not migrate as a sheet, but instead exhibit fascinating and unique patterns of epithelial cell movement, prompting several fundamental questions: 1) How are initial holes in the MES established? This does not require apoptosis, but could instead involve mesenchymal cells pushing through the MES or in a cell-sorting-like behavior driven by differences in cell adhesion or actomyosin contractility between epithelial and mesenchymal cells (Kindberg and Bush, 2019; Lough et al., 2017). Matrix metalloproteinases are induced by TGFβ3 in the MES and required for secondary palate fusion (Blavier et al., 2001), suggesting a role in the initiation of MES breakage. How the positions of these breaks are determined and how local diminution of MES cell-cell adhesion is balanced with overall retention of adherens junctions, which are apparently required for normal collective epithelial migration, is still mysterious. 2) How are MES collectives guided to the oral and nasal surfaces, and do they exhibit cell polarity at the individual, or collective level? We did not observe lamellipodia or other protrusions in MES cell trails, and there was no apparent leading edge to the epithelial trails that were invariably connected to the oral and nasal surface epithelium through a network of trails from the earliest stages. This begs the question of whether and how do epithelial collectives know which direction to migrate? Though islands also did not display directed cellular protrusions, they tended to follow the same path of previous epithelial trails when close to the oral or nasal edge, suggesting that the extracellular matrix (ECM) may act to guide epithelial collectives to the surface. 3) How are cell fate decisions coupled with cell migration behaviors? Based on the migration patterns we observed, MES cells contribute to the nasal palatal surface, which will become respiratory pseudostratified ciliate epithelium, as well as the oral epithelium, which differentiates into oral stratified squamous epithelium. Whether MES cells are specified to these ultimate fates at the time they are undergoing migration, or after they reach their destination may help us to understand how cell migration and fate specification are co-regulated.

Actomyosin contractility driven by NMIIA and enriched filamentous actin at the epithelial/mesenchymal interface is crucial for collective migration. By genetically disrupting non-muscle myosin IIA specifically within the epithelium, we demonstrate an autonomous requirement for actomyosin contractility in collective epithelial migration of the MES. Given the importance of actomyosin contractility in reinforcing cell-cell junctions (Vicente-Manzanares et al., 2009), it is not surprising that loss of NMIIA resulted in a dispersal of MES cells. Dispersed MES cells did not undergo EMT and also did not clear from the palatal mesenchyme, indicating that collective migration involving cell-cell adhesion is crucial. This finding is particularly notable in light of findings that the MYH9 gene, and genes encoding multiple cell-cell adhesion components have been identified to be involved in cleft palate in humans and mice (Birnbaum et al., 2009; Chiquet et al., 2009; Cox et al., 2018; Jia et al., 2010; Lough et al., 2017; Lough et al., 2020; Peng et al., 2016, 9). In addition, strong actomyosin contractions induce cell blebbing by the detachment of the cortical actomyosin cytoskeleton from the plasma membrane (Charras and Paluch, 2008). During apoptosis, caspase cleavage of ROCK I results in hyperactivation of NMII and promotes actomyosin contractility, resulting in membrane blebbing, formation of apoptotic bodies and disruption of nuclear integrity (Coleman et al., 2001; Croft et al., 2005). It is therefore notable that loss of BAX and BAK did not result in the loss of epithelial cell blebbing during MES migration. Cell blebbing is also thought to be involved in various forms of cell motility, including migration through a three-dimensional ECM (Charras and Paluch, 2008). As it has been previously shown that NMII-generated actomyosin contractility results in apoptosis in human embryonic stem cell culture (Chen et al., 2010), our observation that NMIIA-deficient embryos exhibited reduced cleaved caspase 3 staining, raises the possibility that induction of MES apoptosis is a consequence of the high actomyosin contractility that is moving the epithelial trails.

Live imaging of F-actin suggests that pulsatile actomyosin contractility, which is anisotropically enriched at the epithelial/mesenchymal interface and parallel to the direction of epithelial streaming, drives a peristaltic-like movement of epithelial trails. To our knowledge, this form of epithelial collective migration appears distinct from others that have been described in development, such as in the *Drosophila melanogaster* ovary and zebrafish lateral line (Rørth, 2009). Collective migration of neural crest cells in *Xenopus* occurs through supracellular actomyosin contractility at the rear of a migratory cell group which drives intercalation of rear cells and forward movement (Shellard et al., 2018). Though MES migration also appears to occur through the modality of supracellular migration in which the scale of cell behavior is best described at the collective level (Shellard and Mayor, 2019), the precise mechanical drivers are likely different from NCC migration for at least two reasons. First, the rear of MES trails is not enriched for F-actin, and second, it seems unlikely that such localized actomyosin contractility would provide the force needed to push long, thin epithelial trails through the mesenchyme. Greater similarities might be found with cancer cells, in which actomyosin contractility drives the movement of an epithelial collective with stable cell-cell contacts; or cell streaming, in which actomyosin contractility acts independently in each cell of a collective to allow cell rearrangement while maintaining transient cell-cell contacts (Friedl et al., 2012). In the future, localized manipulation of cell-cell adhesion and actomyosin contractility in the MES will help to elucidate the detailed mechanical drivers of this unique form of collective epithelial migration.

## MATERIALS AND METHODS

### Mouse lines

All animal experiments were performed in accordance with the protocols of the University of California, San Francisco Institutional Animal Care and Use Committee. Mice were socially housed under a twelve-hour light-dark cycle with food and water. All alleles used for the experiments herein have been previously described. *Myh9^lox/lox^* (MGI: 4838521) and *Myh10^lox/lox^* (MGI: 4443039) mice have been previously reported (Jacobelli et al., 2010; Ma et al., 2009) and were maintained in a 129/Sv and C57BL/6J mixed genetic background. The following mouse alleles were backcrossed to and maintained on a congenic C57BL/6J genetic background: *Crect* (MGI: 4887352) (Reid et al., 2011), *Bax^lox/lox^* (MGI: 99702) (Knudson et al., 1995), *Bak^-/-^* (MGI: 1097161) (Lindsten et al., 2000), *Rosa26^mTmG/mTmG^* (MGI: 3716464) (Muzumdar et al., 2007), and *Rosa26^nTnG/nTnG^* (MGI: 5504463) (Prigge et al., 2013). For a full description of genetic crosses used to generate embryos; strain background, sex, and stage of embryos; and numbers of embryos analyzed, please refer to **Table S1**. For genotyping, tail biopsies were collected at post-natal day 10 and either sent to Transnetyx or lysed for in-house PCR. For experimental analyses, embryos were harvested at embryonic stages 14.75-17.5, and littermates were used as controls when necessary.

### Immunofluorescence

For cryosection immunofluorescence experiments, whole embryos were fixed in 4% PFA in PBS, dehydrated through a sucrose gradient, embedded in OCT, and frozen in a dry ice/ethanol bath. Blocks were cut to 12 μm sections using an HM550 (Thermo Scientific) or a CM1900 (Leica) cryostat. Sections were then blocked in 5% normal donkey serum (Jackson ImmunoResearch) and 0.1% Triton-X-100 in PBS prior to incubation in primary antibody at 4°C overnight, washed with PBS, and then incubated in secondary antibody at room temperature for 2 hours. Slides were washed with PBS and mounted with Aquamount solution (Lerner laboratories) before imaging. For whole-MES immunofluorescence experiments, embryo heads were fixed in 4% PFA in PBS, embedded in 5% low-melt agarose, and sectioned in room-temperature PBS to 350μm slices using a CT1000S (Leica) vibratome. Sections were washed in PBS, dehydrated through a methanol gradient, bleached in 15% H2O2 in methanol, and rehydrated. Sections were then blocked in 5% normal donkey serum (Jackson ImmunoResearch) and 0.5% Triton-X-100 in PBS prior to incubation in primary antibody at 37°C for 24 hours, washed with PBS, and then incubated in secondary antibody at 37°C overnight. Sections were washed in PBS, dehydrated in a methanol gradient. Finally, tissue sections were cleared through a benzyl alcohol:ss benzyl benzoate (BABB) in methanol gradient (Ahnfelt-Rønne et al., 2007) before imaging. Images were captured using a Zeiss Cell Observer spinning disk confocal microscope or Laser scanning microscope and analyzed using Zeiss Zen software, Imaris software (Bitplane), and/or ImageJ. The following antibodies and dye were used in this study: anti-rat E-cadherin (Invitrogen, 13-1900, 1:300), anti-rabbit cleaved caspase-3 (Cell signaling, 9661, 1:300), anti-goat p63 (R&D, AF1916, 1:300), anti-chicken GFP (abcam, ab4729, 1:1000) and anti-rabbit deltaNp63 (Biolegend, 619001, 1:300). TUNEL staining was performed using In Situ cell death detection kit (Roche, 11684795910) on coronal cryosections of 12 µm thickness. N, nasal surface; O, oral surface.

### RNAscope in situ hybridization

Whole embryos were fixed in 4% PFA in PBS, dehydrated through a sucrose gradient, embedded in OCT, and frozen in a dry ice/ethanol bath. Blocks were cut to 12 μm sections using an HM550 (Thermo Scientific) or a CM1900 (Leica) cryostat. *In situ* hybridization was performed on the 12 μm sections using a *Cre* probe (Advanced Cell Diagnostics, cat# 312281-C3) and RNAscope Multiplex Fluorescent Reagent Kit v2 (Advanced Cell Diagnostics, cat# 323100) according to the manufacturer’s protocol, with the exceptions of excluding antigen retrieval and reducing protease treatment to five minutes. Slides then followed with cryosection immunofluorescence protocol for co-expression analysis.

### Live imaging

The confocal live imaging approach was adapted from our previous work (Kim et al., 2015; Kim et al., 2017), but applied on fresh secondary palate sagittal thick sections. Embryo heads were dissected and the top of the head, calvaria primordia, and lower jaw were removed in ice cold PBS, then the remaining tissue, including the maxillae was embedded in 5% low-melt agarose stored at 37°C. Blocks were sectioned in ice cold DMEM/F12 media to 250μm slices using a Leica CT1000S vibratome. Regions of interest were confirmed by visualization of the endogenous EGFP reporter on the spinning disk confocal microscope. Sections were laid flat in a 35mm No. 0 glass bottom dish (MatTek corporation) and embedded in a mixture of 37°C pre-warmed culture media and agarose, made by adding 3.5% low-melting agarose solution immediately prior to embedding, to make a final concentration (0.6%) to make “live imaging media”. Culture media consisted of 20% fetal bovine serum (FBS), 2 mM L-glutamine, 100 U/mL penicillin, 100 µg/mL streptomycin, 200 µg/mL L-ascorbic acid to Dulbecco’s Modified Eagle Medium (DMEM)/F12 media (Gilbco DMEM/F12 without phenol red). For experiments that visualized F-actin, we added SiR-actin (Spirochrome, CY SC001, 1:5000) just prior to addition of agarose. Sagittal live imaging of the MES was performed using a Zeiss Cell Observer spinning disk confocal microscope equipped with a 37 °C incubation chamber. Time-lapse images were captured with 488 nm and 647 nm laser excitation for 24 hours at the interval of 15mins. For live imaging presented in Figure 3B; Movie 10, the gamma was adjusted to 1.5 in order to visualize low fluorescence epithelial trails without saturating the surface epithelium.

### Data analysis

All Zen (Zeiss) image files were converted into Imaris software (Bitplane) format and subsequently processed using Imaris. Processing included signal intensity adjustments, 3D and/or time cropping and cell tracking. For cell tracking, nuclear EGFP signal was manually segmented to create 3D spots using “spot” tool in Imaris for individual nuclei. Spots were manually selected and tracked in 3D-renderings by every time point. Bleaching correction was used for adjusting the brightness of movies by adding key frames under the video model.

#### Quantification and statistical analysis

For quantification of apoptosis and cell extrusion relating to figures 1 and 5, cell (casp3+, Ecad+, double positive cells) numbers were counted manually on palate sagittal sections immunostained for E-cadherin and cleaved caspase-3 and counterstained with DAPI, regions of interest were selected according to A-P axis.

For quantification of E-cadherin occupancy relating to figure 2 and 4, images of whole palates were divided transversely into four equal parts by the grids. In this setup, the rostral-most region contains the nasal surface, the next two regions are entirely seam, and the caudal-most region contains the oral surface. E-cadherin signal was segmented using Imaris to generate 3D membrane surfaces in each region, and the volumes of surfaces were measured. Statistical analysis was performed using GraphPad Prism 8. Unpaired t-tests were used to determine statistical significance.

## ACKNOWLEDGEMENTS

We thank Scott Oakes for sharing *Bax* and *Bak* mutant mice. We are grateful to Seungil Kim for initiating breeding of *Bax;Bak* mice and to Fang Shiuan Leung for outstanding genotyping support. We are grateful to our laboratory and UCSF colleagues for helpful conversations and suggestions and to Ace Lewis for critical reading of the manuscript. This work was supported by R01DE025877 and R01DE023337 from NIH/NIDCR to J.O.B.

## FIGURE LEGENDS

**Figure S1.**
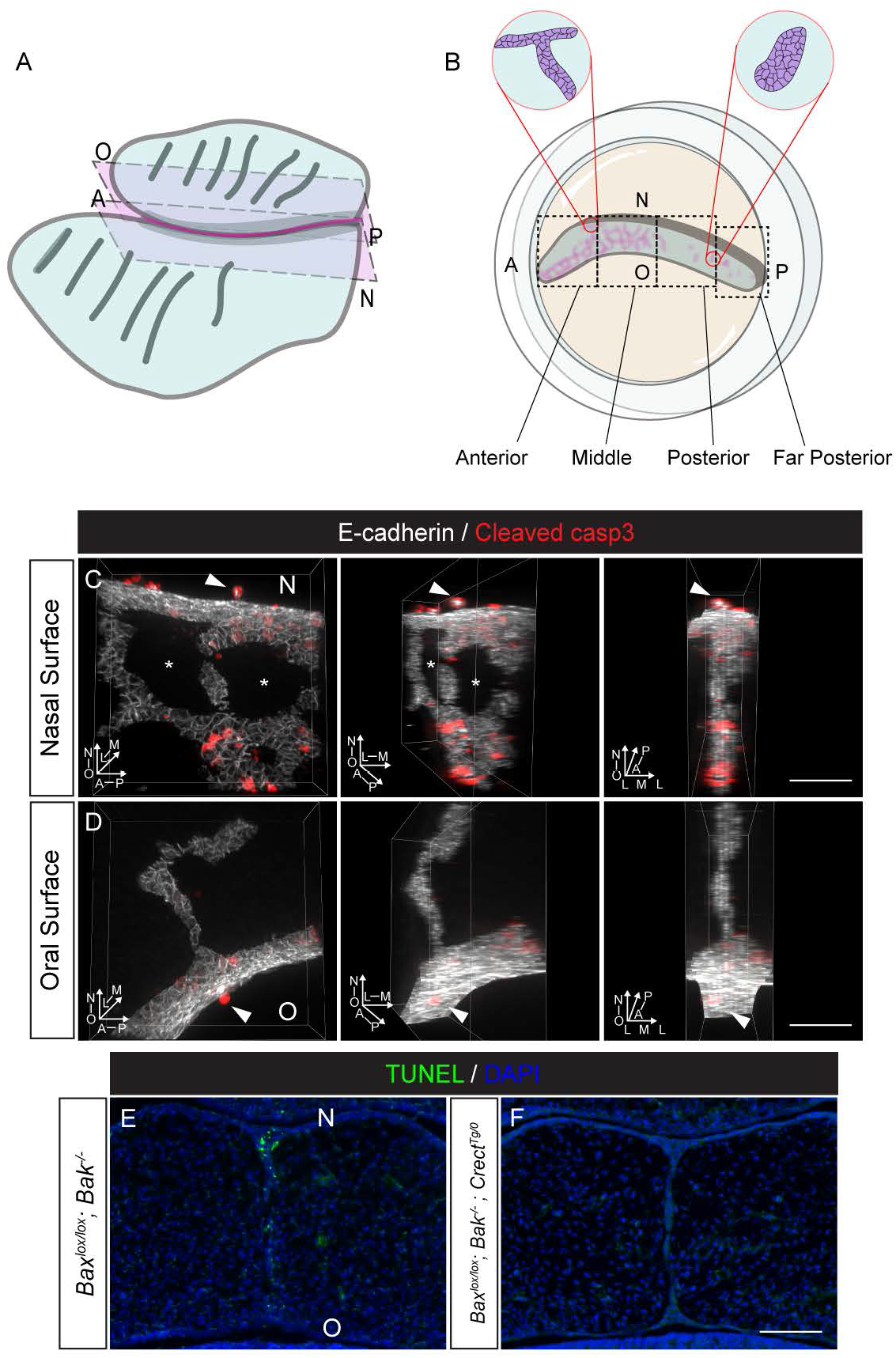
Sagittal section live imaging approach. (A,B) Schematic representation of the sagittal section live imaging approach shows an isolated mouse secondary palate with the oral surface facing up. (A) A secondary palate trimmed using a vibratome to retain a thick section containing the MES region represented between the two dash-lined pink rectangles. (B) The thick section is placed in a glass bottom dish and mounted in 5% low-melt agarose (orange). Views of an epithelial (magenta) trail and an island surrounded by mesenchyme (blue) are magnified in red circles. Black-dashed squares represent the different regions described throughout this paper. (C-D) 3D-renderings of WT sagittal sections are shown focusing at the region including the nasal surface (C) and oral surface (D). Sequential panels show the rendering turned at 45°and 90° on the nasal-oral axis. Sections were immunostained for E-cadherin (white) and cleaved caspase-3 (red) to reveal MES cells and apoptotic MES cells. Arrowheads indicate extruded MES cells on nasal and oral surfaces. Asterisks point to holes in epithelium. N, nasal surface; O, oral surface; L, lateral; M, medial; A, anterior; P, posterior. Scale bar, 50μm. (E) TUNEL staining reveals an abundance of cell death in coronal sections of control *Bax^lox/lox^ Bak^-/-^* secondary palatal shelves, (F) Loss of cell death from the MES of *Bax^lox/lox^; Bak^-/-^ Crect^Tg/0^* mutant embryos. Scale bar, 50 μm

**Figure S2.**
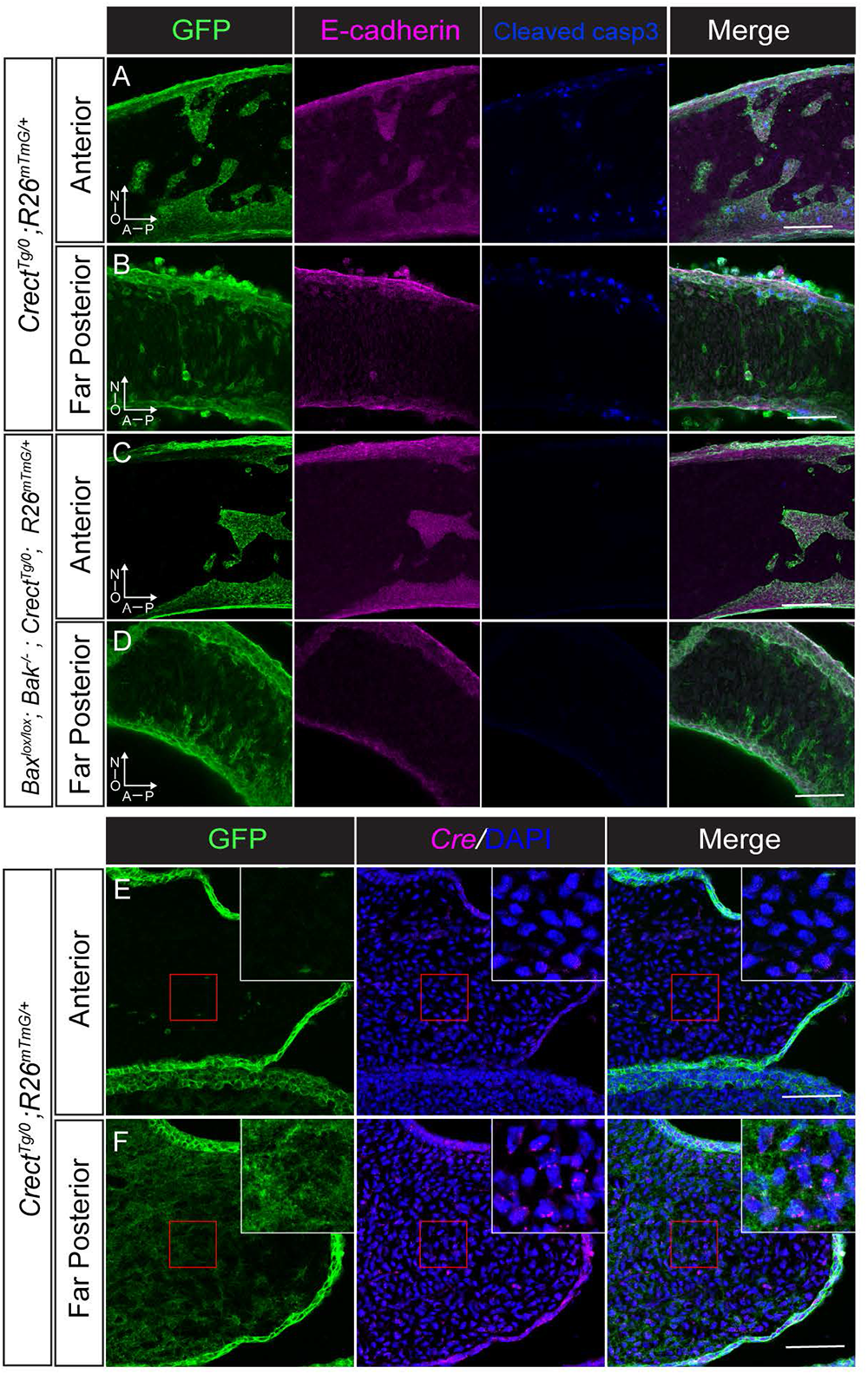
EMT does not compensate for BAX/BAK loss in MES removal. (A-D) Epithelial lineage tracing and immunostaining of GFP (green) and E-cadherin (magenta) on sagittal thick sections reveals no GFP-expressing, E-cadherin negative cells in the anterior palate of control (A) *Crect^Tg/0^; R26^mTmG/+^* or (C) *Bax^lox/lox^; Bak^-/-^; Crect^Tg/0^; R26^mTmG/+^* mutant embryos. An abundance of GFP-expressing, E-cadherin negative cells were observed in the posterior palate of control (C) *Crect^Tg/0^; R26^mTmG/+^* and (D) *Bax^lox/lox^; Bak^-/-^; Crect^Tg/0^; R26^mTmG/+^* mutant embryos. Scale Bar, 100μm. N, nasal surface; O, oral surface; L, lateral; M, medial; A, anterior; P, posterior (E-F) Detection of *Cre* RNA expression (magenta) by RNAScope *in-situ* hybridization relative to *Crect^Tg/0^; R26^mTmG/+^* recombination in the anterior (E) and far posterior (F) palatal shelves of *Crect^Tg/0^; R26^mTmG/+^* embryos. Inset is a high magnification view of the red square. Scale bars, 100μm.

**Figure S3.**
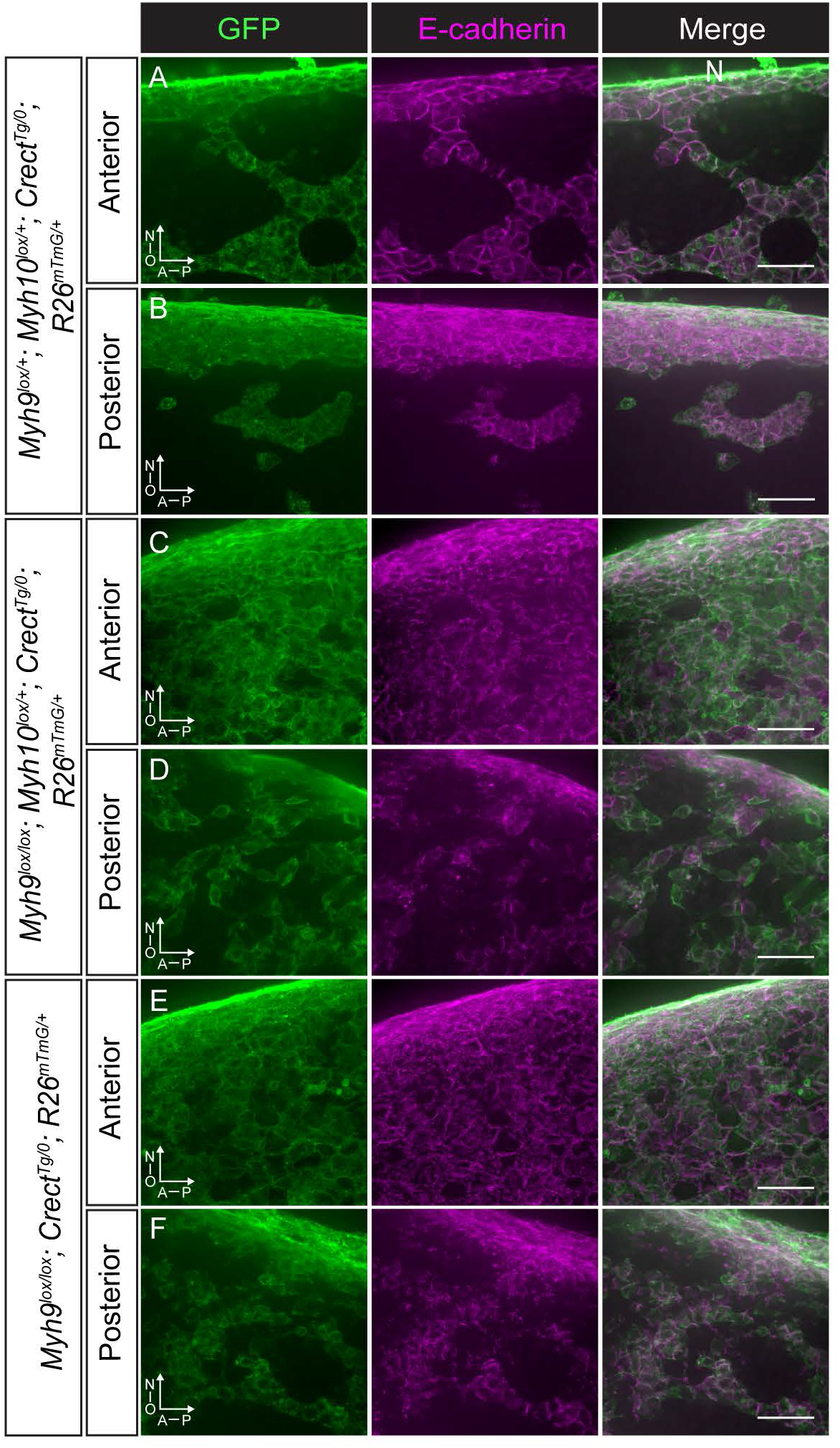
NMIIA and not NMIIB is required for collective organization of the MES. 3D-rendered images of E15.5 control and NMII mutants palates visualizing *Crect*-mediated recombination (GFP in green) and MES cells, as labeled by E-cadherin in magenta. (A,B) In *Myh9^lox/+^; Myh10^lox/+^; Crect^Tg/0^; R26^mTmG/+^* controls, trails and islands are detected in both anterior (A) and posterior (B) palate regions, with cell-cell junctions clearly revealed by E-cadherin expression. (C,D) In *Myh9^lox/lox^; Myh10^lox/+^; Crect^Tg/0^; R26^mTmG/+^* and (E, F) *Myh9^lox/lox^; Crect^Tg/0^; R26^mTmG/+^* mutants, a large number of MES cells are broadly maintained at the seam in the anterior palate (C,E, respectively) and diffusely distributed in the posterior palate (D,F, respectively). Moreover, cellular distribution of E-cadherin is disrupted in mutant MES cells. Scale bar, 25μm.

**Figure S4.**
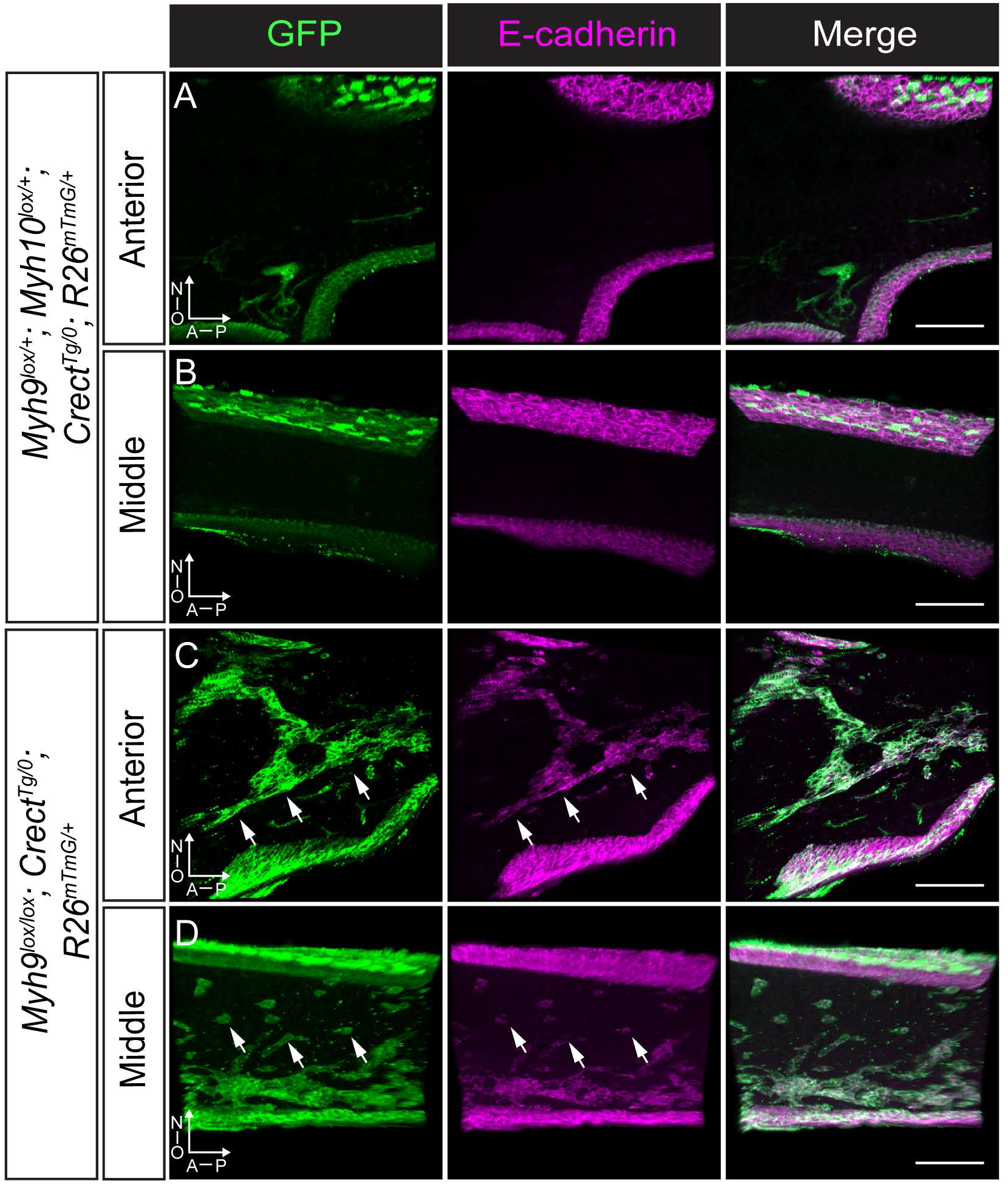
Retention of epithelial inclusions in secondary palate upon loss of NMIIA. 3D-rendered images of secondary palate thick sagittal sections immunostained for eGFP (green), to label *Crect*-mediated recombination, and E-cadherin (magenta), to label MES cells. (A,B) No MES cells are detected in *Myh9^lox/+^; Myh10^lox/+^; Crect^Tg/0^; R26^mTmG/+^* control secondary palate at E17.5 (C,D) Persistent MES cells are detected in the mutant *Myh9^lox/lox^; Crect^Tg/0^; R26^mTmG/+^* palate (white arrows point to some examples). A, anterior; P, posterior; N, nasal surface; O, oral surface. Scale Bar, 100μm.

---

**Movie 1. Live imaging of the initiation of MES breakdown during secondary palate fusion.** Confocal live imaging of GFP expression in *Crect^Tg/0^; R26^mTmG/+^* secondary palate sagittal section at E14.75 shows breakage of the MES in an early stage of MES removal. Red arrowheads indicate membrane blebbing. High magnification images of Z20 (3μm interval) were captured every 15 minutes for 20 hours. Scale Bar, 25μm.

**Movie 2. Organization of MES epithelial cells into MES trails** Confocal live imaging of GFP expression in *Crect^Tg/0^; R26^mTmG/+^* secondary palate sagittal section at E15.5 shows MES trail movement and breakage. Arrowheads point to trails and broken ends of a trail. Renderings are performed by Imaris with high magnification imaging of Z20 (3μm interval). Images were captured every 15mins for 20 hours. Scale Bar, 50μm.

**Movie 3. The MES migrates in collective epithelial trails** MES cells in a trail exhibit collective migration. GFP expressing cells from an E15.25 *Crect^Tg/0^; R26^nTnG/+^* secondary palate were live imaged and manually tracked. Tracking shows all MES cells in the trail following the same direction during migration. Each colored sphere and corresponding line indicate a single cell and its trajectory. Images were captured every 15 minutes for 5.75 hours. Scale Bar, 40μm.

**Movie 4. MES islands located near the surface epithelium are removed by cell migration.** Confocal live imaging of GFP expression in *Crect^Tg/0^; R26^mTmG/+^* secondary palate sagittal section at E15.5 shows an island moving towards and merge into the nearby oral surface. Renderings are performed by Imaris with high magnification imaging of Z20 (3μm interval). Images were captured every 15mins for 20 hours. Scale Bar, 50μm.

**Movie 5. MES islands located far from epithelial surfaces are removed by apoptosis.** Confocal live imaging of GFP expression in *Crect^Tg/0^; R26^mTmG/+^* secondary palate sagittal section at E15.5 shows an island that is far from an epithelial surface exhibit stereotyped apoptotic appearance and progressive shrinkage of the epithelial island. Arrowheads point out characteristic blebbing behavior of apoptotic cells. Renderings are performed by Imaris with high magnification imaging of Z20 (3μm interval). Images were captured every 15mins for 20 hours. Scale Bar, 40μm.

**Movie 6 Loss of apoptosis does not disrupt MES breakage.** MES sheet breakage in a *Bax; Bak* mutant at an early stage of MES removal. Confocal live imaging of *Bax^lox/lox^; Bak^-/-^; R26^mTmG/+^; Crect^Tg/0^* secondary palate sagittal section at E14.75 shows the breakage of MES. Red arrowheads are pointing to membrane blebbing. Renderings were performed in Imaris with high magnification imaging of Z20 (3μm interval). Images were captured every 15mins for 20 hours. Scale Bar, 25μm.

**Movie 7. Loss of apoptosis does not disrupt collective epithelial migration.** Confocal live imaging of GFP expression in *Bax^lox/lox^; Bak^-/-^; R26^mTmG/+^; Crect^Tg/0^* secondary palate sagittal section at E15.5 focusing on trails shows collective MES cell movement. Arrowheads indicate trails in the movie. Renderings are performed by Imaris with high magnification imaging of Z20 (3μm interval). The gamma setting for this movie is adjusted to 1.5. Images were captured every 15 minutes for 20 hours. Scale Bar, 50μm.

**Movie 8. MES epithelial islands do not shrink or disappear upon loss of apoptosis.** Confocal live imaging of GFP expression in *Bax^lox/lox^; Bak^-/-^; R26^mTmG/+^; Crect^Tg/0^* secondary palate sagittal section at E15.5 focusing on an island shows the lack of MES cell removal by apoptosis. Arrowheads are pointing to membrane blebbing that still occurs in mutants. Renderings are performed by Imaris with high magnification imaging of Z20 (3μm interval). Images were captured every 15 minutes for 20 hours. Scale Bar, 40μm.

**Movie 9. MES removal through collective epithelial migration in control embryos.** Confocal live imaging of GFP expression in control *Myh9^lox/+^; Myh10^lox/+^; R26^mTmG/+^; Crect^Tg/0^* secondary palate sagittal section at E15.5 shows normal MES cells movement in the trail. Renderings are performed by Imaris with high magnification imaging of Z20 (3μm interval). Images were captured every 15 minutes for 20 hours. Scale Bar, 40μm.

**Movie 10. Collective organization and migration is disrupted upon loss of NMIIA.** Confocal live imaging of GFP expressing cells in *Myh9^lox/lox^; R26^mTmG/+^; Crect^Tg/0^* secondary palate sagittal section at E15.5 shows lack of directional MES cell movement. Renderings are performed in Imaris with high magnification imaging of Z20 (3μm interval). Images were captured every 15 minutes for 20 hours. Scale Bar, 40μm.

**Movie 11. Anisotropic actin accumulation in epithelial trails drives collective epithelial movement.** Actin contracting contributes to MES cell migration. (Relating to Fig. 6E) Secondary palate sagittal section of E15.5 *Crect^Tg/0^; R26^mTmG/+^* embryos were stained with SiR-actin and then live imaged for GFP expressing MES cells and F-actin. Arrowheads indicate contracting F-actin that enable MES cell movement. Scale Bar, 50μm. (Relating to Fig. 6F) A magnified view focusing on the trail reveals contractile actin filament adjacent to MES cells of the trail. Scale Bar, 10μm. Renderings are performed by Imaris with high magnification imaging of Z5 (3μm interval). Images were captured every 15mins for 6 hours (E) or 2.5 hours (F).

